# *C. elegans* CLASP/CLS-2 negatively regulates membrane ingression throughout the oocyte cortex and is required for polar body extrusion

**DOI:** 10.1101/2020.04.02.021675

**Authors:** Aleesa J. Schlientz, Bruce Bowerman

## Abstract

The requirements for oocyte meiotic cytokinesis during polar body extrusion are not well understood. In particular, the relationship between the oocyte meiotic spindle and polar body contractile ring dynamics remains largely unknown. We have used live cell imaging and spindle assembly defective mutants lacking the function of CLASP/CLS-2, kinesin-12/KLP-18, or katanin/MEI-1 to investigate the relationship between meiotic spindle structure and polar body extrusion in *C. elegans* oocytes. We show that spindle bipolarity and chromosome segregation are not required for polar body contractile ring formation and chromosome extrusion in *klp-18* mutants, but oocytes with severe spindle assembly defects due to loss of CLS-2 or MEI-1 have penetrant and distinct polar body extrusion defects: CLS-2 is required early for contractile ring assembly or stability, while MEI-1 is required later for contractile ring constriction. We also show that CLS-2 negatively regulates membrane ingression throughout the oocyte cortex during meiosis I, and we explore the relationship between global cortical dynamics and oocyte meiotic cytokinesis.

**Author Summary:** The precursor cells that produce gametes—sperm and eggs in animals—have two copies of each chromosome, one from each parent. These precursors undergo specialized cell divisions that leave each gamete with only one copy of each chromosome; defects that produce incorrect chromosome number cause severe developmental abnormalities. In oocytes, these cell divisions are highly asymmetric, with extra chromosomes discarded into small membrane bound polar bodies, leaving one chromosome set within the much larger oocyte. How oocytes assemble the contractile apparatus that pinches off polar bodies remains poorly understood. To better understand this process, we have used the nematode *Caenorhabditis elegans* to investigate the relationship between the bipolar structure that separates oocyte chromosomes, called the spindle, and assembly of the contractile apparatus that pinches off polar bodies. We used a comparative approach, examining this relationship in three spindle assembly defective mutants. Bipolar spindle assembly and chromosome separation were not required for polar body extrusion, as it occurred normally in mutants lacking a protein called KLP-18. However, mutants lacking the protein CLS-2 failed to assemble the contractile apparatus, while mutants lacking the protein MEI-1 assembled a contractile apparatus that failed to fully constrict. We also found that CLS-2 down-regulates membrane ingression throughout the oocyte surface, and we explored the relationship between oocyte membrane dynamics and polar body extrusion.

## Introduction

Oocyte meiosis comprises a single round of genome replication followed by two highly asymmetric cell divisions that produce a single haploid gamete and two small polar bodies that contain discarded chromosomes (1–4). An acentrosomal spindle segregates homologous chromosomes during the first reductional division, called meiosis I, and half of the recombined homologs are extruded into the first polar body. The equational meiosis II division then segregates sister chromatids, with half extruded into a second polar body and half remaining in the oocyte cytoplasm. Despite being essential for reducing oocyte ploidy, little is known about the cues that organize and influence the actomyosin contractile ring that mediates polar body extrusion.

The oocyte contractile ring initially forms distal to the membrane-proximal meiotic spindle pole, with the spindle axis oriented orthogonally to the overlying cell cortex (5). During anaphase the contractile ring ingresses past both the membrane-proximal pole and one set of the segregating chromosomes to then constrict and ultimately separate the nascent polar body from the oocyte. These dynamics contrast substantially with mitotic cytokinesis (Fig 1A), during which signals from astral microtubules and the central spindle position the contractile ring midway between the two spindle poles (6–8). While the signals required for contractile ring assembly and constriction during mitotic cytokinesis are relatively well understood, how oocyte meiotic spindles influence contractile ring dynamics during polar body extrusion is not known.

**Fig 1.**
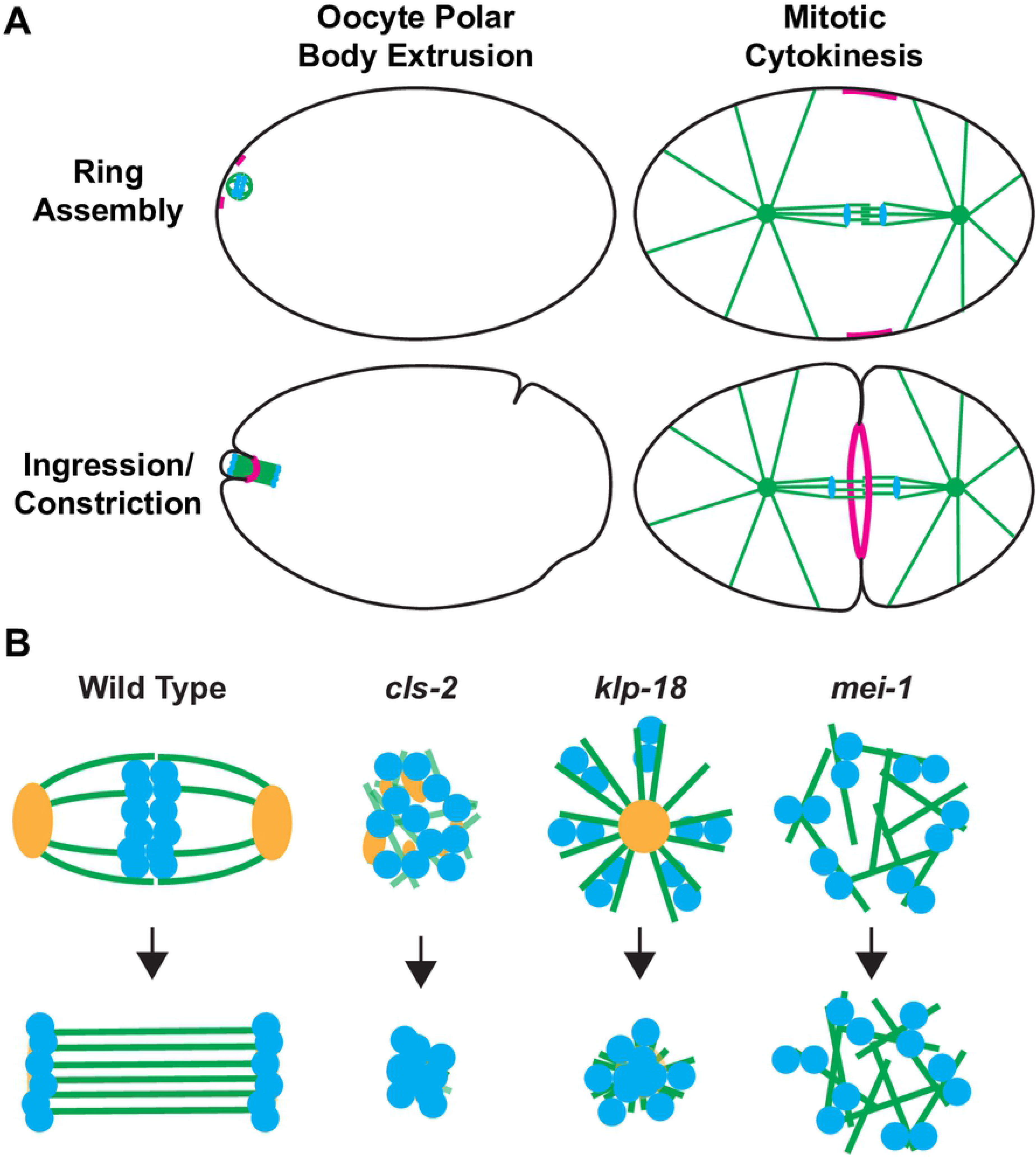
Schematics of oocyte meiotic polar body extrusion, mitotic cytokinesis and oocyte meiotic spindle assembly-defective mutants. (A) The positioning and dynamics of contractile ring assembly and ingression during oocyte polar body extrusion and mitotic cytokinesis. (B) Illustrations of oocyte meiotic spindle structure in control and mutant oocytes. Green = microtubules, blue = chromosomes, magenta = contractile rings, and orange = ASPM-1 pole marker, black = plasma membrane. See text for details.

In *Caenorhabditis elegans*, several genes are required for polar body extrusion, but how their functions are coordinated remains poorly understood. Similar to mitotic cytokinesis, polar body cytokinesis requires filamentous actin and the non-muscle myosin II heavy-chain NMY-2 and light-chains MLC-4 and MLC-5 (9–11). The cytoskeletal scaffolding protein anillin/ANI-1 facilitates transformation of the initial actomyosin contractile ring into a midbody tube, with anillin depletion resulting in large and unstable polar bodies that often fuse with the oocyte (12). Consistent with its role as a key activator of cortical actomyosin, the small GTPase RhoA (RHO-1) and its RhoGEF ECT-2 also are required for oocyte polar body extrusion (10, 13). Knockdown of the centralspindlin complex, comprised of MgcRacGAP/CYK-4 and kinesin-6/ZEN-4, results in the assembly of abnormally large contractile rings and a subsequent failure in extrusion (10). Finally, the chromosomal passenger complex (CPC) member Aurora B/AIR-2 also is required (14).

Formation of the contractile ring distal to both meiotic spindle poles raises the question of how the ring moves relative to the spindle such that it constricts midway between segregating chromosomes. One mechanism proposed for *C. elegans* is that global contraction of actomyosin throughout the oocyte cortex produces a hydrostatic cytoplasmic force that, combined with depletion of cortical actomyosin overlying the membrane-proximal pole, leads to an out-pocketing of the membrane within the contractile ring and pushes the spindle into the protruding pocket (10, 15). In support of this hypothesis, increased global cortical contractility due to depletion of casein kinase 1 gamma (CSNK-1), a negative regulator of RhoA activity, often results in extrusion of the entire meiotic spindle (15). However, assessing whether global cortical contractility is required for polar body extrusion has been challenging due to the overlap in requirements for global cortical contractility and polar body contractile ring assembly and constriction.

Despite a stereotyped spatial relationship between the oocyte meiotic spindle and contractile ring assembly and ingression, little is known about how the spindle might influence ring assembly and dynamics. Nevertheless, four observations suggest that the meiotic spindle provides important cues. First, meiotic spindles that fail to translocate to the oocyte cortex induce the formation of membrane furrows that ingress deeply towards the displaced spindle (10). Second, loss of the centralspindlin complex, which localizes to the central spindle during anaphase, results in the formation of rings with an abnormally large diameter (10). Third katanin/*mei-1* mutants, which assemble apolar spindles, produce very large polar bodies (16, 17). Finally, while work in mice suggests that chromosomes themselves may provide cues for ring assembly and polar body extrusion via the small GTPase Ran (18, 19), knock down of *C. elegans* RAN-1 does not prevent polar body extrusion (20, 21). These findings suggest that in *C. elegans* the oocyte meiotic spindle provides cues that influence contractile ring assembly and ingression.

To explore the relationship between meiotic spindle assembly and polar body extrusion, we have examined oocyte cortical actomyosin dynamics in three spindle assembly defective mutants that each lack the function of a conserved protein: CLASP/CLS-2, kinesin-12/KLP-18, or katanin/MEI-1 (Fig 1B). CLASP family proteins promote microtubule stability through their association with microtubules and their tubulin heterodimer-binding TOG (Tumor Over-expressed Gene) domains, decreasing the frequency of microtubule catastrophe and promoting rescue of de-polymerizing microtubules (22–25). Moreover, human CLASPs have been shown to influence not only microtubule stability and dynamics, but also to interact with actin filaments and potentially crosslink filamentous actin and microtubules (26). CLS-2 is one of three *C. elegans* CLASPs and is required for mitotic central spindle stability as well as oocyte meiotic spindle assembly and chromosome segregation (27–30). Vertebrate kinesin-12/KLP-18 family members promote mitotic spindle bipolarity by contributing to forces that push apart anti-parallel microtubules (31–33). Consistent with such a function, *C. elegans* oocytes lacking the kinesin-12 family member KLP-18 form monopolar meiotic spindles that draw chromosomes towards a single spindle pole and thus fail in chromosome segregation (34–36). The widely conserved microtubule severing complex katanin is encoded by two *C. elegans* genes, *mei-1* and *mei-2* (37, 38). Loss of either subunit results in the formation of apolar meiotic spindles that fail to congress or segregate chromosomes (34, 39, 40).

Here we report our use of fluorescent protein fusions and live cell imaging to characterize polar body extrusion during meiosis I in mutants lacking the function of CLS-2, KLP-18 or MEI-1. Previous studies indicate that both CLS-2 and MEI-1 are involved in polar body extrusion, with oocytes lacking CLS-2 frequently failing to extrude polar bodies (27, 30, 41), and oocytes lacking MEI-1 forming very large polar bodies (16, 17, 42, 43). Furthermore, *klp-18* mutants sometimes lack an oocyte pronucleus, suggesting that all oocyte chromosomes can be extruded (34). However, the process of polar body extrusion has not been directly examined in any of these mutants. Our live imaging of contractile ring assembly and ingression in these mutant backgrounds shows that bipolar spindle assembly and chromosome segregation are not required for oocyte contractile ring assembly and polar body extrusion. However, CLS-2 is required for proper contractile ring assembly or stability and also acts as a negative regulator of global cortical membrane ingressions, while MEI-1 may be required late in polar body extrusion for contractile ring constriction.

## Results

### CLS-2 is required for oocyte meiotic spindle assembly and polar body extrusion

To investigate the role of CLS-2, we first examined the localization of a CLS-2∷GFP fusion in live oocytes (Fig 2A, S1 Fig, S1 and S2 Movies). Consistent with previous reports, CLS-2∷GFP initially localized to meiosis I spindle microtubules and kinetochore cups, and to small patches dispersed throughout the oocyte cortex (27, 30, 41). Around the time of anaphase onset, the cortical CLS-2∷GFP patches disappeared and CLS-2∷GFP localized to the central spindle between the segregating chromosomes (27, 30, 41). These results suggest that CLS-2 might have roles not only at the oocyte meiotic spindle but also throughout the oocyte cortex.

**Fig 2.**
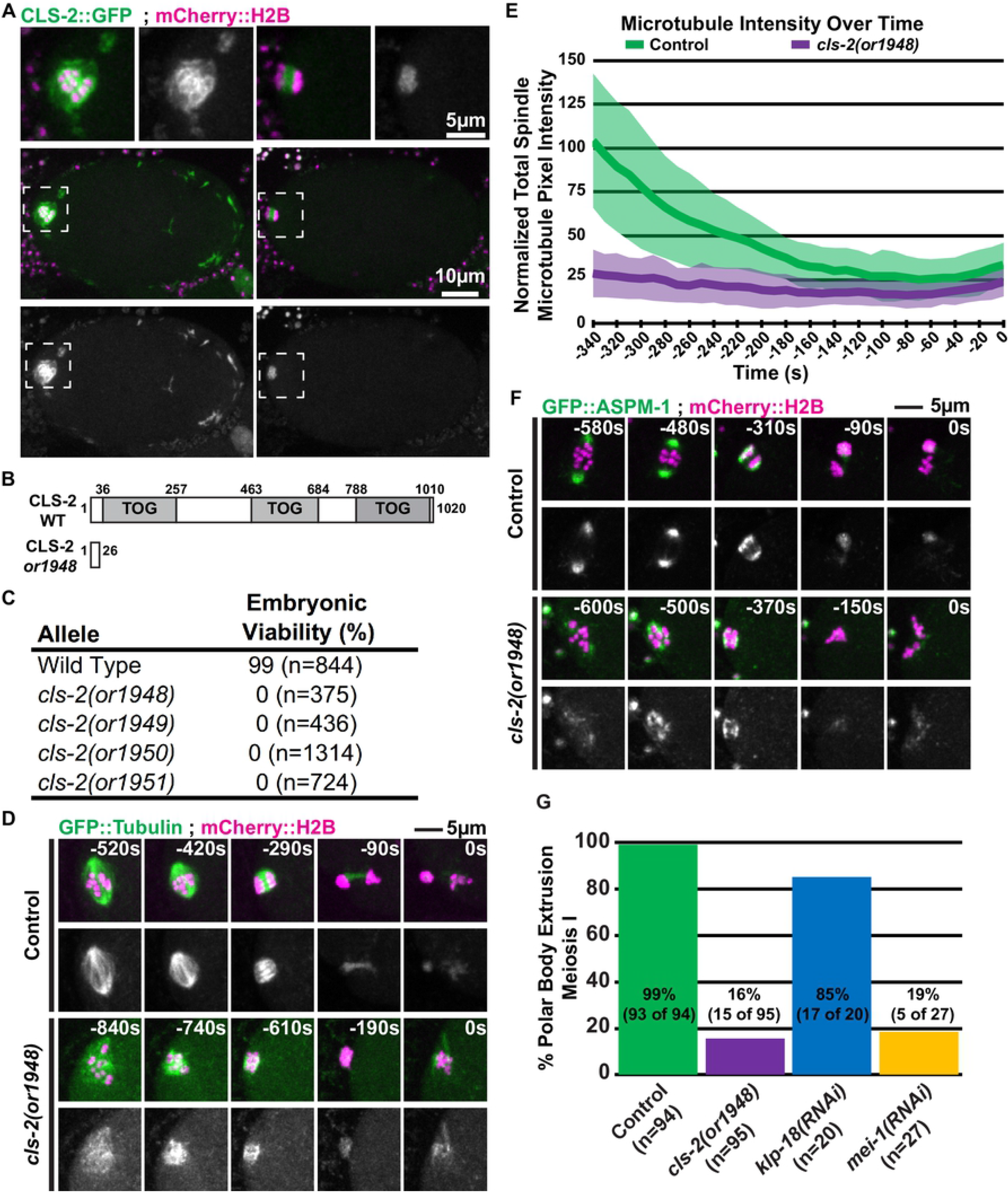
CLS-2 is required for oocyte meiotic spindle assembly and polar body extrusion. (A) Control oocytes expressing CLS-2∷GFP and mCherry:H2B during meiosis I; dashed boxes indicate the zoomed-in regions shown in top row. (B) Protein domain map for wild type CLS-2 (28), and location of first premature stop codon due to frameshift in *cls-2(or1948)*. (C) Table of embryonic viability in wild type (N2) and *cls-2* CRISPR-generated loss of function alleles. (D) *C*ontrol and mutant oocytes expressing GFP∷Tubulin and mCherry∷H2B during meiosis I; t = 0 seconds (0s) corresponds to the end of meiosis I and beginning of meiosis II (see Materials and methods). (E) Comparison of integrated spindle microtubule pixel intensity over time between control (n = 17) and *cls-2* mutant oocytes (n = 9 @ −340s, n = 10 @ −330 to −300s, n = 11 @ - 290 to −160s, and n = 12 @ −150 to 0s), with the average intensity indicated and one standard deviation indicated. Here and in subsequent figure panels, T = 0s corresponds to the end of meiosis I and beginning of meiosis II, unless indicated otherwise (see Materials and methods). (F) *C*ontrol and mutant oocytes expressing GFP∷ASPM-1 and mCherry∷H2B during meiosis I. (G) Percent of control and mutant oocytes that extrude a polar body during meiosis I, as determined by mCherry or GFP-fused histone signal remaining stably ejected from the oocyte cytoplasm for the duration of imaging (see Materials and methods).

Previous studies of CLS-2 requirements have used RNA interference (RNAi) or auxin-induced degradation of degron-tagged CLS-2 to reduce its function and have emphasized its central spindle function. To more definitively assess its roles during oocyte meiosis, we used CRISPR/Cas9 to generate putative null alleles. Each of the four alleles we isolated contains small insertions or deletions that result in frame shifts and premature stop codons before the first TOG domain, likely making them null (Fig 2B, S1 Fig). All are recessive, and homozygous mutant hermaphrodites exhibit fully penetrant embryonic lethality (Fig 2C). To investigate CLS-2 requirements, we have used the *cls-2(or1948)* allele, and hereafter we refer to oocytes from homozygous *cls-2(or1948)* hermaphrodites as *cls-2* mutants.

We first used *ex utero* live cell imaging with transgenic strains that express GFP fused to a β-tubulin (GFP∷TBB-2) and mCherry fused to a histone H2B (mCherry∷H2B) to examine microtubule and chromosome dynamics during meiosis I in control and *cls-2* mutant oocytes (Fig 2D; see Materials and methods). Control oocytes formed barrel shaped bipolar spindles that shortened and rotated to become perpendicular to the oocyte cortex prior to anaphase and polar body extrusion (n=19). Consistent with previous reports (27, 30, 41), the meiosis I spindles in *cls-2* mutants were disorganized and lacked any obvious bipolarity, with chromosomes moving into a small cluster before failing to segregate (n=13). Furthermore, microtubule levels appeared to be reduced, and quantification of the integrated spindle microtubule intensity over time showed a substantial reduction in microtubule levels throughout meiosis I compared to control oocytes (Fig 2E), consistent with the established roles of CLASP family members in promoting microtubule stability (see Introduction). To assess when defects in meiosis I spindle assembly first appear in *cls-2* oocytes, we used *in utero* live cell imaging and observed the early assembly of a normal cage-like microtubule structure that surrounded the oocyte chromosomes, followed by a rapid collapse of this microtubule structure to form an abnormally small cluster associated with the oocyte chromosomes (S1 Fig) (n=5). To further examine spindle assembly in *cls-2* mutants, we also imaged meiosis I in oocytes from transgenic strains expressing an endogenous fusion of GFP to the pole marker ASPM-1 and mCherry∷H2B (Fig 2F). As described previously (44), multiple small GFP∷ASPM-1 foci coalesced to form a bipolar spindle in control oocytes (n=14). In *cls-2* mutants, the GFP∷ASPM-1 foci failed to coalesce to form two spindle poles but rather moved over time to form a single tight cluster of multiple small foci, and chromosomes again moved into a small cluster and failed to segregate (n=11). We conclude that CLS-2 plays an important role in promoting microtubule stability throughout meiosis I and is required early in bipolar spindle assembly and for chromosome segregation.

In addition to the extensive meiotic spindle defects, we also observed that *cls-2* mutants frequently failed to extrude a polar body at the end of meiosis I, consistent with previous reports (27, 30). Control oocytes regularly extruded a polar body at the end of meiosis I, as scored using transgenic strains expressing either GFP∷H2B or mCherry∷H2B and observing whether oocyte chromosomes remained extruded for the duration of live imaging (93 of 94 oocytes; Fig 2G). In contrast, polar body extrusion failed in about 84% of the *cls-2* mutant oocytes (80 of 95, Fig 2G). This finding suggests that CLS-2 might play a direct role in polar body extrusion, or that the spindle defects observed in *cls-2* mutants might indirectly disrupt extrusion.

### Polar body extrusion defects in *cls-2* mutants are not due to absence of the central spindle-associated proteins AuroraB/AIR-2 or MgcRacGAP/CYK-4

Because *cls-2* mutant oocytes fail to assemble a central spindle or segregate chromosomes and CLS-2∷GFP localizes to the central spindle, and because the central spindle-associated protein AuroraB/AIR-2 and the centralspindlin component MgcRacGAP/CYK-4 are required for polar body extrusion (10, 14, 27, 45), we first considered whether the polar body extrusion defects might be due to a failure to localize central spindle proteins to the disorganized spindle microtubules in *cls-2* oocytes. To address this possibility, we used transgenic strains expressing GFP fusions to either AIR-2 or CYK-4 and examined their localization in control and *cls-2* mutant oocytes. As reported previously, both proteins localized to the central spindle during anaphase in control oocytes (Figs 3A and 3B) (n=12 GFP∷AIR-2 and n=12 CYK-4∷GFP). In *cls-2* mutants, despite the lack of spindle organization, both AIR-2 and CYK-4 localized to the spindle region (Figs 3A and 3B) (n=15 GFP∷AIR-2 and n=11 CYK-4∷GFP). While we cannot rule out the possibility that improper localization of AIR-2 and CYK-4 to the defective spindle might be responsible for the failure to extrude polar bodies in *cls-2* mutants, this failure cannot be explained by a simple loss of these central spindle-associated proteins from the mutant spindle microtubules.

**Fig 3.**
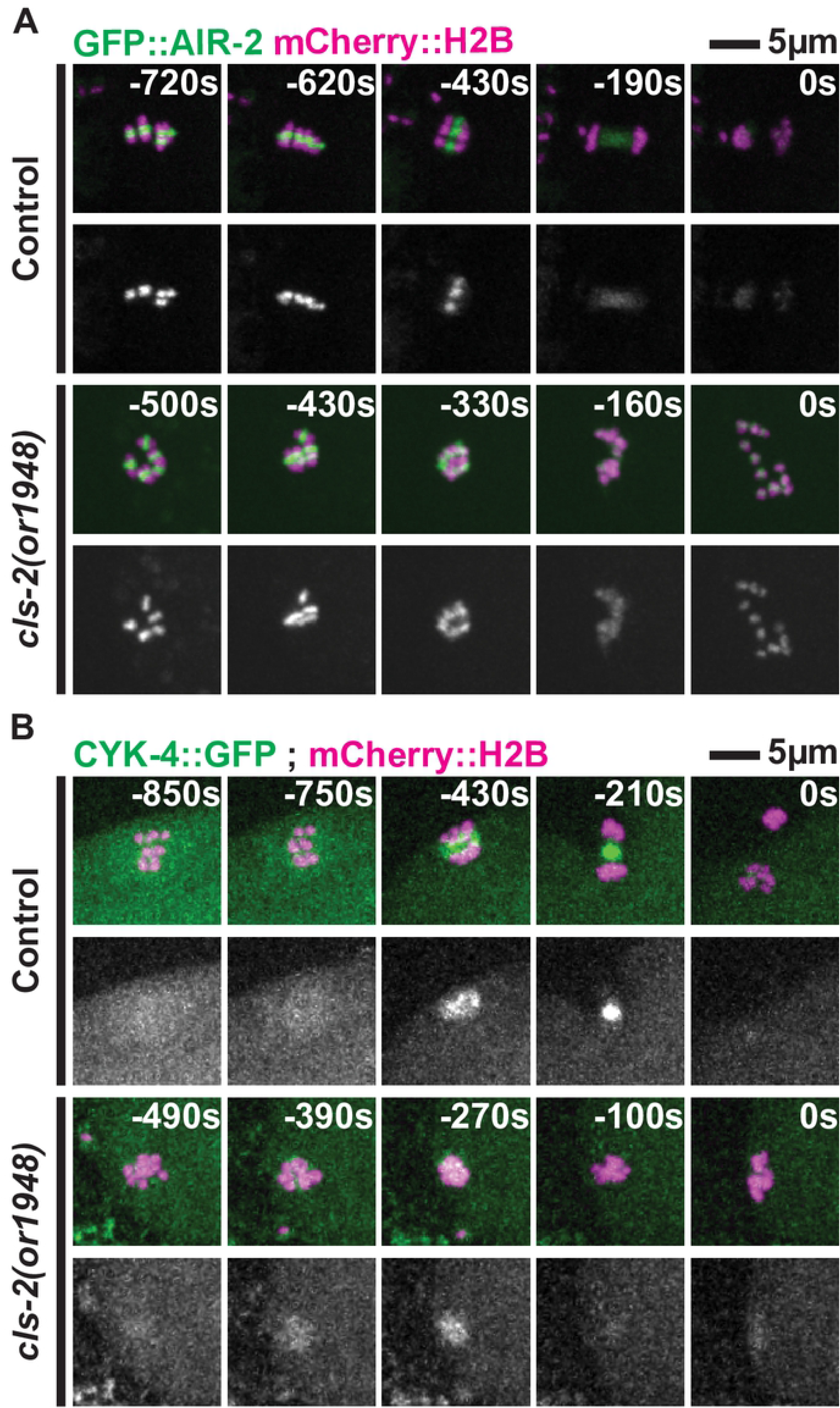
Central spindle proteins AIR-2 and CYK-4 still associate with spindle microtubules in *cls-2* mutant oocytes. Control and *cls-2* mutant oocytes expressing GFP∷AIR-2 and mCherry∷H2B (A) or GFP∷CYK-4 and mCherry∷H2B (B) during meiosis I.

### CLS-2 and MEI-1, but not spindle bipolarity, are required for polar body extrusion

Because the relationship between spindle structure and polar body extrusion is unclear, we next took a comparative approach and also examined meiosis I polar body extrusion after using RNAi to knock down either kinesin-12/KLP-18 or katanin/MEI-1. As illustrated schematically in Fig 1B, *klp-18* mutants assemble monopolar spindles while *mei-1* spindles are apolar, and both fail to segregate chromosomes (see Introduction). We first simply assessed whether polar body extrusion was successful, again using live imaging with transgenic strains expressing either GFP∷H2B or mCherry∷H2B fusions and scoring whether chromosomes were retained in polar bodies for the duration of imaging (Fig 2G). In *klp-18* mutants, chromosomes were successfully retained in a polar body in 15 of 20 oocytes. In contrast, after MEI-1 knockdown, chromosomes were extruded and retained in a polar body in only 5 of 27 oocytes. The absence of meiosis I polar body extrusion in *mei-1* mutant oocytes was surprising because *mei-1* mutants were originally described as typically having abnormally large polar bodies (16, 17), but how often polar body extrusion succeeds or fails has not been reported. Our data indicate that meiosis I polar body extrusion usually fails and suggest that the abnormally large polar bodies observed in *mei-1* mutants result from defects in meiosis II (see Discussion).

Based on the frequent success of polar body extrusion in *klp-18* mutants, we conclude that spindle bipolarity and chromosome segregation are not required for polar body extrusion. Thus, the failed extrusions in *cls-2* and *mei-1* mutants are not simply due to an absence of spindle bipolarity or a failure to segregate chromosomes.

### Meiotic spindle-associated furrows are abnormal in *cls-2 and mei-1* mutant oocytes

To better understand the polar body extrusion defects in oocytes lacking CLS-2 or MEI-1, we next examined the spindle-associated membrane furrows that mediate polar body extrusion in transgenic strains expressing mCherry fused to a plasma membrane marker (mCherry∷PH) and GFP∷H2B (Figs 4A and 4B, S2 Fig, S3 and S4 Movies). In 9 of 16 control oocytes, we observed early membrane furrows that ingressed until they pinched together to encapsulate and extrude chromosomes into the first polar body. In 6 of 16 control oocytes we observed early membrane furrows that were not as clearly resolved in our imaging data but eventually led to polar body extrusion, and in one oocyte furrows were not obvious but a polar body was nevertheless extruded.

**Fig 4.**
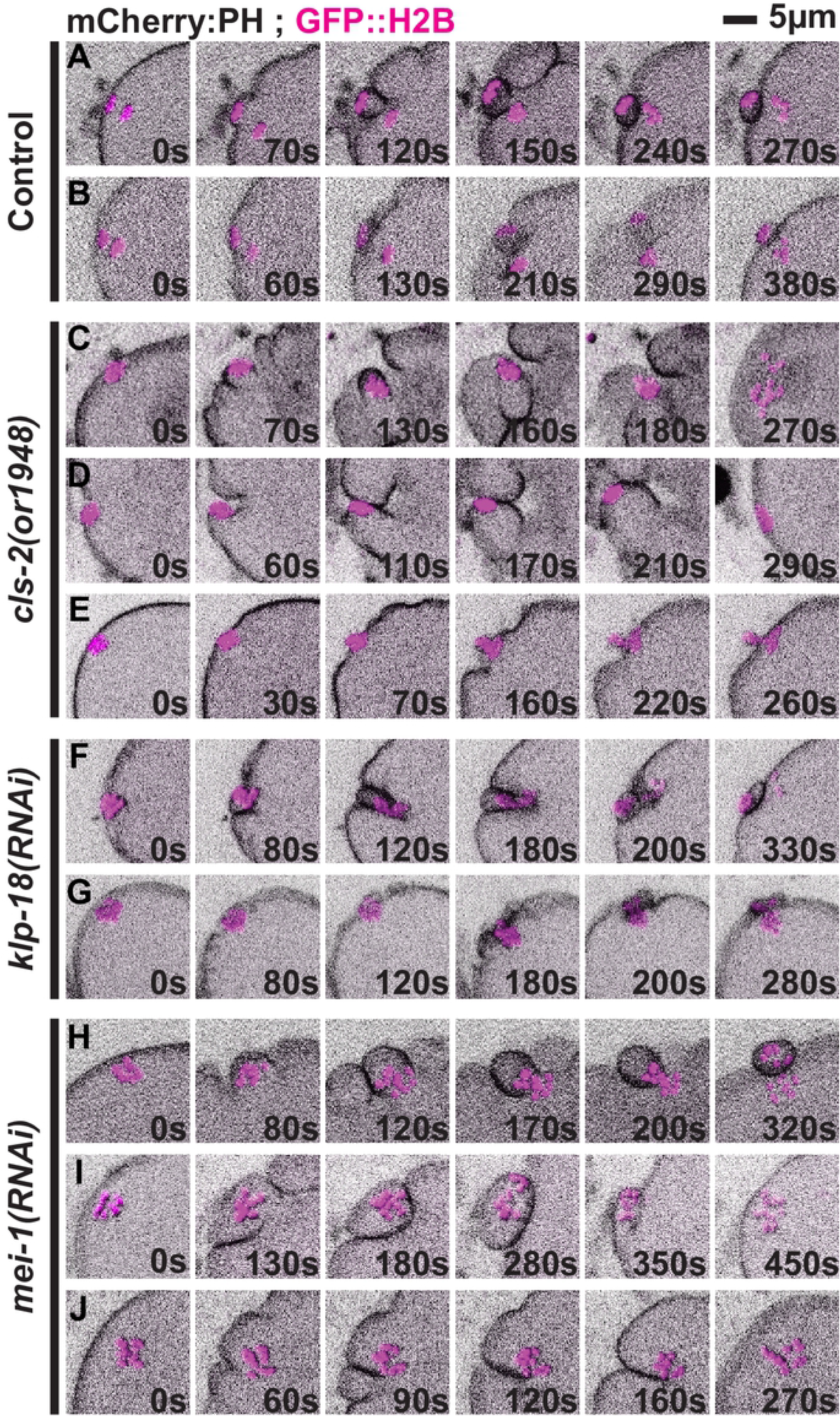
Spindle-associated membrane furrowing during meiosis I in control and mutant oocytes. Images show projections of five focal planes encompassing the chromosomes during meiosis I in oocytes expressing an mCherry fusion to a PH domain to mark the plasma membrane using sum projections and GFP∷H2B using maximum projections. Representative examples of control (A, B), *cls-2* mutant (C-E), *klp-18(RNAi)* (F, G) and *mei-1(RNAi)* (H, I) oocytes. Polar body extrusion failed in C-E and H, and was successful in all others. T = 0s corresponds to the timepoint immediately before cortical furrowing begins. See text for details.

In contrast, we observed extensive spindle-associated membrane furrowing defects in *cls-2* mutants (Figs 4C-E, S3 and S4 Figs, S5 and S6 Movies). In 7 of 19 oocytes we observed two furrows in cross-section that retracted before pinching together, one oocyte that formed two furrows that pinched together but failed late in polar body extrusion, and one oocyte that formed two furrows and successfully extruded a polar body (S3 Fig). One of 19 oocytes appeared to form three spindle-associated furrows in cross-section and extruded a polar body (S3 Fig). In 7 of 19 oocytes we observed only a single visible spindle-associated furrow in cross-section that ingressed either to one side of, or directly toward the oocyte chromosomes (Fig 4D, S4 Fig), suggesting that the contractile ring collapsed into a more linear ingressing structure rather than maintaining a ring-like shape. Finally, in one oocyte the membrane dynamics were indistinct, but chromosomes were extruded into a polar body (Fig 4E), and in one oocyte there was no obvious spindle-associated furrowing and polar body extrusion failed (S4 Fig).

In oocytes depleted of *klp-18*, we observed furrows that more nearly resembled those in control oocytes (Figs 4F and G, S5 Fig, S7 and S8 Movies). In 4 of 10 oocytes, we observed two furrows in cross-section that ingressed more deeply and then pinched together (Fig 4F), and only 1 of these 4 failed in polar body extrusion. In 6 of 10 oocytes, we observed shallow furrows adjacent to the oocyte chromosomes (Fig 4G), and only 1 of these 6 failed to extrude a polar body.

After MEI-1 knockdown, we observed furrows that initially resembled those in control oocytes but were more widely spaced and often failed late during constriction (Figs 4H-J, S6 Fig, S9 and S10 Movies). In 2 of 11 oocytes, we observed two furrows in cross-section that ingressed and pinched together to extrude a polar body (Fig 4H). In 3 of 11 oocytes two furrows ingressed and pinched together but then regressed and released chromosomes back into the oocyte cytoplasm (Fig 4I). In 5 of 11 oocytes two furrows ingressed but retracted before pinching together and failed in polar body extrusion (Fig 4J), and finally in 1 of 11 oocytes we observed only a single spindle-associated furrow in cross-section that failed to extrude a polar body.

To summarize, in *klp-18* mutant oocytes we observed spindle-associated furrows that usually encapsulated chromosomes and extruded polar bodies, although the oocyte chromosomes were often in close proximity to the membrane with furrows that were shallow and difficult to detect. In *cls-2* and *mei-1* mutants, meiosis 1 polar body extrusion frequently failed but we observed distinct defects. While membrane furrowing initially appeared normal but eventually failed in most *mei-1* mutant oocytes, *cls-2* oocytes often appeared abnormal early in furrow ingression and exhibited more severe defects as extrusion attempts progressed.

### Polar body contractile ring dynamics are more severely defective in the absence of CLS-2 than in *klp-18* or *mei-1* mutant oocytes

We next examined assembly and ingression of the contractile ring during oocyte meiosis I, using live cell imaging with transgenic strains expressing both a GFP fusion to the non-muscle myosin II NMY-2 and mCherry∷H2B. In 11 of 11 control oocytes, NMY-2∷GFP foci initially assembled into discontinuous but discernible rings over the membrane proximal pole, after spindle rotation and before extensive chromosome segregation, and then became more continuous and prominent as they ingressed and constricted between the segregating chromosomes to extrude polar bodies (Fig 5A, S7 Fig, S11 and S12 Movies).

**Fig 5.**
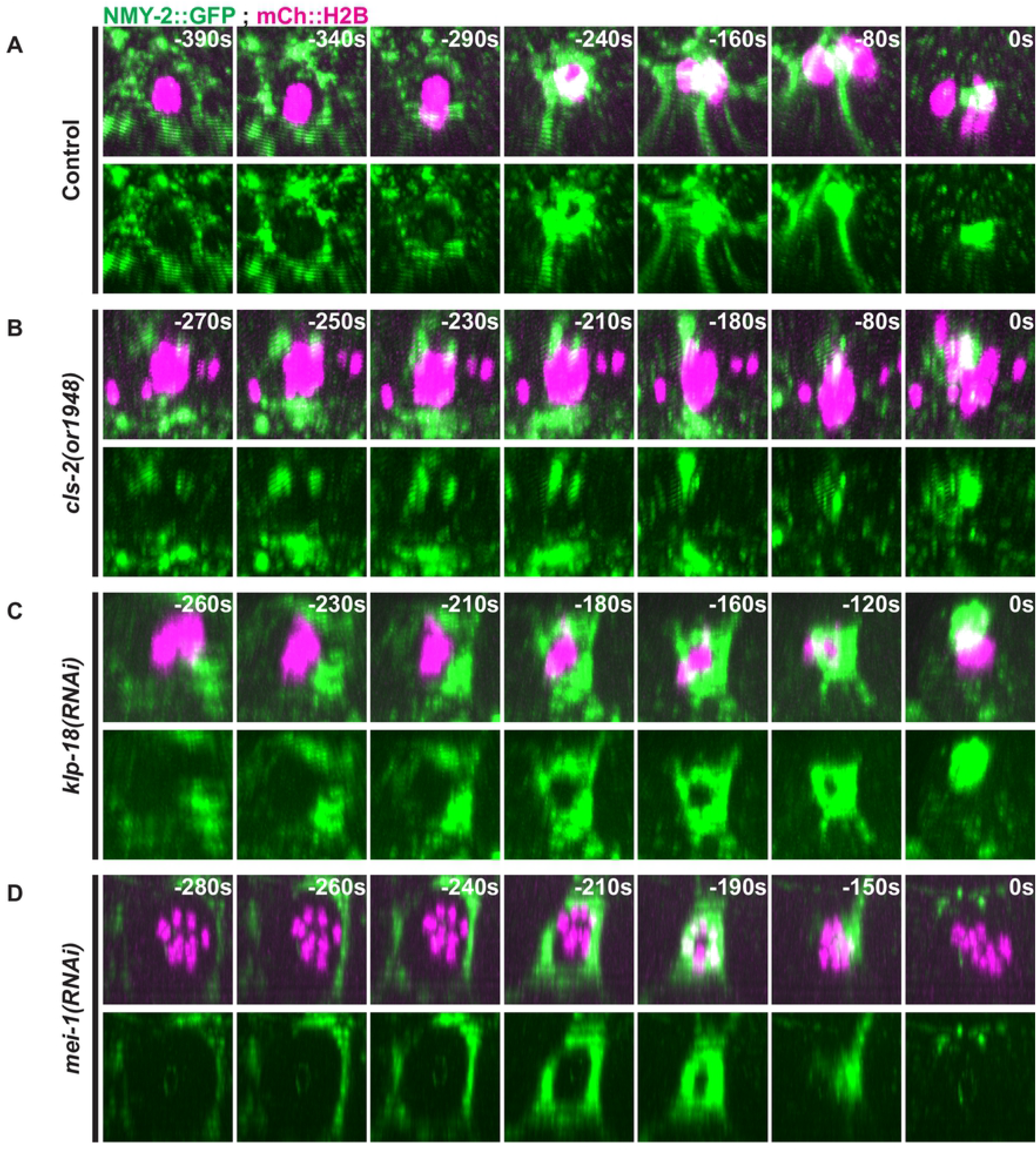
Contractile ring non-muscle myosin NMY-2 dynamics during meiosis I in control and mutant oocytes. Three-dimensionally projected and rotated images of control (A), *cls-2* mutant (B), *klp-18(RNAi)* (C), and *mei-1(RNAi)* (D) oocytes expressing NMY-2∷GFP and mCherry∷H2B. Z-stacks were rotated as to look down on contractile ring assembly and dynamics over time.

In *cls-2* mutant oocytes, the assembly and stability of NYM-2∷GFP contractile ring structures were severely defective (Fig 5B, S8 Fig, S13 and S14 Movies). In 7 of 11 oocytes, fragmented or partial contractile rings assembled and 6 of these oocytes failed to extrude a polar body, while 3 of 11 oocytes formed abnormal assemblies of NMY2∷GFP that were more linear and not ring-like, although 2 of these extruded a polar body. Finally, 1 of 11 oocytes formed a relatively normal looking contractile ring that extruded a polar body. The fragmented or partial contractile rings observed in *cls-2* mutants often collapsed into single bright foci or bands during constriction.

Contractile ring assembly and dynamics appeared much more normal in *klp-18* mutant oocytes (Fig 5C, S9 Fig, S15 and S16 Movies). In 10 of 10 oocytes after KLP-18 knockdown, NMY-2∷GFP foci assembled into rings that ingressed and constricted with dynamics similar to those observed in control oocytes, and in 9 of the 10 oocytes chromosomes were stably extruded into polar bodies.

In MEI-1 knockdown oocytes, ring assembly and ingression were much more normal compared *to cls-2* mutants, but we nevertheless observed a range of later defects and eventual failures to extrude polar bodies (Fig 5D, S10 Fig, S17 and S18 Movies). In 11 of 13 oocytes, the NMY-2∷GFP rings that initially formed were larger in diameter compared to control oocytes, and in 3 of these 11 oocytes the rings constricted and successfully extruded a polar body. In another 5 of these 11 oocytes, the rings constricted extensively but ultimately regressed and failed at polar body extrusion, while in 3 the rings ingressed and only constricted partially before regressing and failing to extrude polar bodies. Finally, in 2 of 13 oocytes, ring assembly and ingression were more defective and polar body extrusion failed.

To further characterize the polar body extrusion defects in *cls-2* mutants, we also examined ring assembly and dynamics in transgenic strains expressing mNeonGreen fused to the anillin ANI-1 (mNG∷ANI-1), which is dispensable for assembly of the actomyosin contractile ring but required for its conversion from a ring to a tube during constriction (46), and mCherry∷H2B (Fig 6A, S19 and S20 Movies). In 10 of 10 control oocytes, mNG∷ANI-1 assembled into contractile rings that ingressed and constricted between segregating chromosomes to extrude a polar body, while in 10 of 10 *cls-2* oocytes a fragmented contractile ring structure formed and failed to extrude a polar body. We also used two-color live imaging to examine NMY-2∷mKate2 and mNG∷ANI-1 simultaneously, and observed that these two contractile ring components were co-localized in both control and *cls-*2 mutant oocytes (Fig 6B, S21 and S22 Movies) (n=11 control, n=13 *cls-2(or1948)*). We conclude that CLS-2 is required for proper ring assembly and ingression dynamics of not only NMY-2 but also ANI-1.

**Fig 6.**
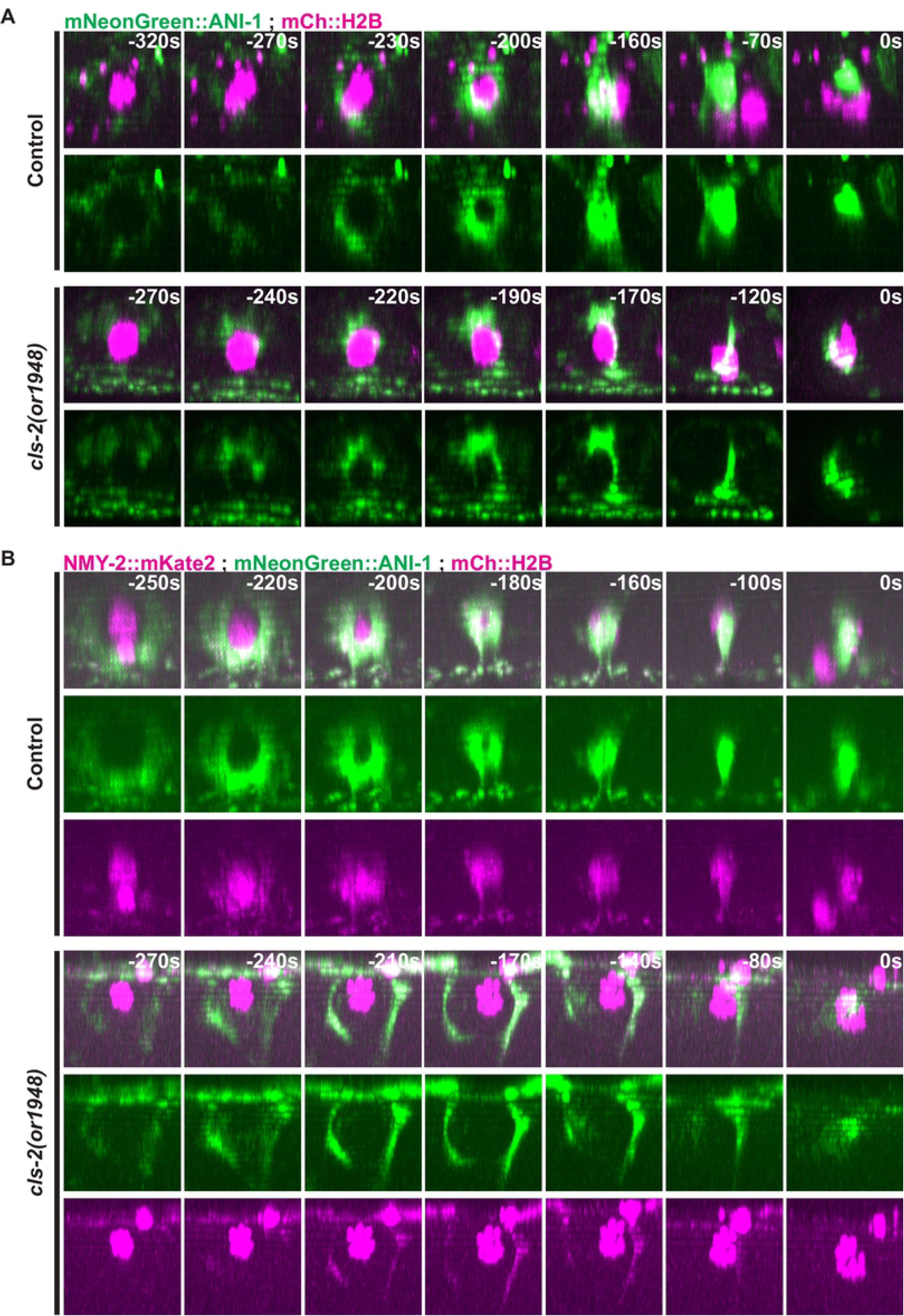
Contractile ring anillin ANI-1 and non-muscle myosin NMY-2 dynamics during meiosis I in control and *cls-2* mutant oocytes. Three-dimensionally projected and rotated images of control and *cls-2* mutant oocytes expressing mNeonGreen∷ANI-1 and mCherry∷H2B (A) or NMY-2∷mKate2, mNeonGreen∷ANI-1, and mCherry∷H2B, with overlays shown in top row for each set. Z-stacks were rotated so as to look down on contractile ring assembly and dynamics over time.

### CLS-2 negatively regulates membrane ingression throughout the oocyte cortex during meiosis I

Global contraction of the oocyte actomyosin cortex has been proposed to promote polar body extrusion by generating a hydrostatic cytoplasmic force that (i) produces an out-pocketing of the actomyosin depleted membrane inside the meiotic contractile ring, and (ii) pushes the spindle partially into the extruded membrane pocket (see Introduction). To explore the relationship between spindle structure, global cortical contractility and polar body extrusion, we next examined membrane ingressions throughout the oocyte cortex during meiosis I in control and mutant oocytes, using transgenic strains expressing mCherry∷PH and GFP∷H2B fusions. To document these membrane ingressions, we used temporal overlays of a single central z-plane to portray simultaneously the membrane position at all time points throughout the period of global cortical furrowing. In control oocytes, we observed the spindle-associated furrows and a small number of additional furrows along the oocyte cortex (Fig 7A, S11 Fig) (n=16).

**Fig 7.**
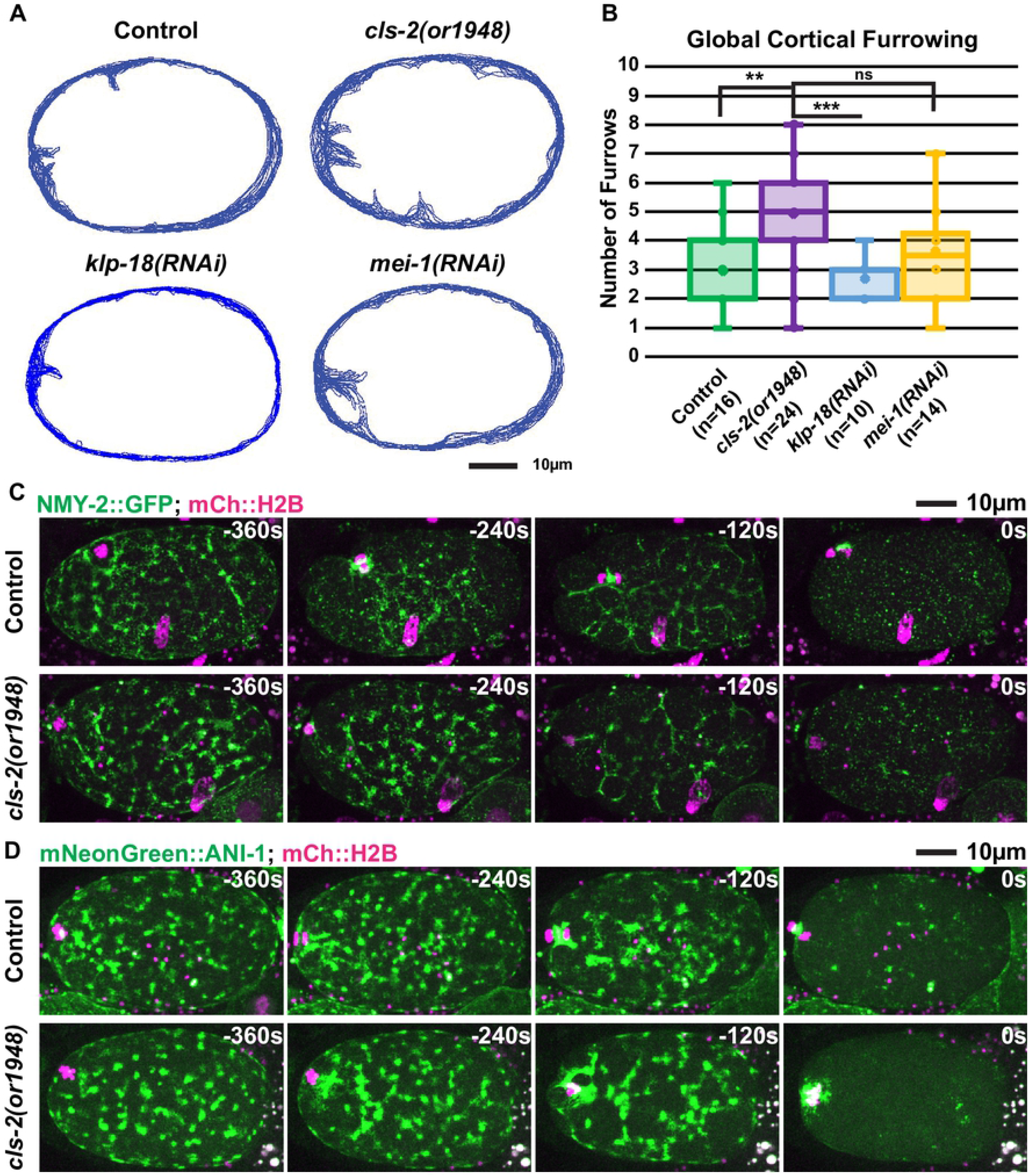
Membrane ingressions throughout the oocyte cortex during meiosis I in control and mutant oocytes. (A) Membrane temporal overlays for control, *cls-2* mutant, *klp-18(RNAi)*, and *mei-1(RNAi)* oocytes representing the membrane positions of a single focal plane during the period of meiosis I global cortical furrowing for a single oocyte of each genotype. (B) Quantification of the number of global cortical furrows in control and mutant oocytes (see Materials and methods). t-Test results: Control vs *cls-2(or1948)* p = 0.0014 (**), *cls-2(or1948)* vs *klp-18(RNAi)* p = 3.14E-5 (***), *cls-2(or1948)* vs *mei-1(RNAi)* p = 0.054 (ns). (C and D) *C*ontrol and *cls-2* mutant oocytes expressing (C) NMY-2∷GFP and mCherry∷H2B or (D) mNeonGreen∷ANI-1 and mCherry∷H2B.

In contrast, *cls-2* mutant oocytes exhibited more extensive cortical furrowing compared to control oocytes (n=24), while oocytes depleted of either KLP-18 or MEI-1 more nearly resembled control oocytes (Fig 7A, S12 and S13 Figs, S3-10 Movies) (n=10 *klp-18(RNAi)* and n=14 *mei-1(RNAi)*). Quantification of global cortical furrowing showed that *cls-2* oocytes had significantly more furrows compared to control and *klp-18* oocytes (Fig 7B). We conclude that CLS-2 negatively regulates global cortical furrowing, and we suspect that the CLS-2∷GFP patches detected throughout the oocyte cortex are responsible for this regulation (Fig 2A, S1 Fig, S1 and S2 Movies; see Discussion).

We next examined the dynamics of the cortical actomyosin cytoskeleton, which mediates the furrowing that occurs throughout the oocyte cortex during polar body extrusion. In control oocytes, NMY-2 and ANI-1 localized to dynamic patches throughout the oocyte cortex during meiosis I contractile ring assembly and ingression, and then dissipated late in anaphase when global cortical furrowing ends (Figs 7C and 7D, S14 and S15 Figs, S23 and S24 Movies) (n=11 NMY-2∷GFP, n=10 mNG∷ANI-1). To determine if the increased global cortical furrowing in *cls-2* oocytes is caused by an increase in NMY-2 or ANI-1 patch size or duration, we examined the dynamics of NMY-2∷GFP and mNG∷ANI-1 and observed dynamics similar to those in control oocytes (Figs 7C and 7D, S16-18 Figs, S25 and S26 Movies) (n=11 NMY-2∷GFP, n=10 mNG∷ANI-1), and we did not detect any statistically significant difference in the area occupied by the cortical NMY-2∷GFP patches throughout the period of global cortical furrowing and polar body extrusion. These data suggest that the excess global cortical furrowing observed in *cls-2* oocytes is not due to altered NMY-2 or ANI-1 patch dynamics. While the increased global cortical furrowing in CSNK-1 knockdown oocytes is associated with altered NMY-2 and ANI-1 cortical patch dynamics and with extrusion of the entire meiosis I spindle into polar bodies (15), the increased global cortical furrowing in *cls-2* mutants does not correlate with altered cortical patch dynamics and is instead associated with polar body extrusion failure. Thus, the relationship between global cortical furrowing and polar body extrusion is complex and requires further investigation (see Discussion).

## Discussion

Remarkably little is known about the relationship between spindle structure and contractile ring assembly and constriction during oocyte meiotic cell division. To gain insight into the cues that influence contractile ring dynamics during polar body extrusion, we have examined contractile ring assembly and constriction in three different *C. elegans* spindle assembly-defective mutants. Our results indicate that the *C. elegans* CLASP family member CLS-2 is required not only for assembly of a bipolar meiosis I spindle and for chromosome segregation, but also for an early stage in oocyte meiosis I contractile ring assembly or stability. This requirement for CLS-2 is not due simply to a failure in assembling bipolar spindles or to a lack of chromosome segregation, because *klp-18/kinesin-12* mutant oocytes formed monopolar spindles and failed to segregate chromosomes but did assemble stable contractile rings that usually extruded chromosomes into a polar body. We have further shown that *mei-1/katanin* mutant oocytes, which assemble apolar and disorganized oocyte meiosis I spindles, also usually failed to extrude polar bodies. However, while CLS-2 was required early for contractile ring assembly or stability, MEI-1 appears to be required to limit the initial diameter of the contractile ring and for later steps in ring constriction. Finally, we also observed increased global cortical furrowing during meiosis I in *cls-2* mutant oocytes but not in *klp-18* or *mei-1* mutants, raising the possibility that proper regulation of cortical membrane ingressions throughout the oocyte may be required for contractile ring assembly or stability during polar body cytokinesis. Our results highlight the value of exploring the relationship between mutant oocyte meiotic spindle structures and polar body contractile ring dynamics.

### Using spindle assembly defective mutants to explore contractile ring dynamics during oocyte polar body extrusion

While the requirements for several factors that control oocyte meiotic spindle assembly in *C. elegans* have been described (1–3), the impact of the resulting spindle assembly defects on polar body extrusion has remained largely unexplored. Indeed, anecdotal observations have suggested that polar body extrusion can occur even in mutants with severe oocyte spindle assembly defects. For example, reducing the function of the microtubule severing complex katanin, comprised of MEI-1 and −2 in *C. elegans*, results in severely defective apolar spindles that have greatly reduced levels of microtubules and fail to organize or segregate chromosomes and yet often produce abnormally large polar bodies. However, the dynamics of contractile ring assembly, ingression and constriction in *mei-1/katanin* mutants has not been directly examined. Similarly, mutant oocytes lacking the kinesin-12 family member KLP-18 assemble monopolar spindles that fail to segregate chromosomes but often produce zygotes that completely lack an egg pronucleus, indicating that all of the oocyte chromosomes are sometimes extruded into polar bodies, but again the process of polar body extrusion in *klp-18* mutants has not been directly examined. In mouse oocytes, DNA-coated beads have been shown to promote the assembly of a polar body-like cortical protrusion, which requires the small GTPase Ran that mediates chromatin signaling and, if allowed to assemble a bead-associated spindle structure, can lead to successful extrusion (18, 19). However, knockdown of the *C. elegans* Ran family member RAN-1 does not prevent chromosome segregation or polar body extrusion (20, 21), suggesting that chromatin cues do not mediate contractile ring assembly. Moreover, meiotic spindles that fail to translocate to the oocyte cortex induce the formation of membrane furrows that ingress deeply toward the spindle (10), and *C. elegans* mutant oocytes lacking central spindle proteins have been shown to produce abnormally large contractile rings that often fail to extrude chromosomes into a polar body (10), suggesting that the oocyte meiotic spindle does influence ring assembly and function.

Given the critical roles of astral and central spindle microtubules in promoting the assembly of a contractile ring during mitosis (6–8), the distinct dynamics and geometry of oocyte meiotic contractile rings compared to mitotic contractile rings (Fig 1A), and the indications that oocyte meiotic spindles influence contractile ring dynamics in *C. elegans,* we have characterized polar body extrusion in *C. elegans* mutants that are severely defective in oocyte spindle assembly. We were first motivated to do so based on our observation that *cls-2* oocytes frequently failed to extrude chromosomes, in contrast to anecdotal and indirect observations that other *C. elegans* mutants with severe oocyte meiotic spindle assembly defects can extrude chromosomes into polar bodies.

Because *cls-2* mutant oocytes failed to assemble bipolar spindles or segregate chromosomes, we also examined polar body extrusion in *klp-18/kinesin-12* mutant oocytes that assemble monopolar spindles and also fail to segregate chromosomes (34–36). In contrast to *cls-2* mutants, in which contractile ring assembly or stability and membrane ingression were severely defective, contractile ring assembly and ingression appeared much more normal in *klp-18* mutant oocytes and chromosomes were usually extruded into a polar body. We conclude that the defects in *cls-2* mutant oocytes are not due a failure to assemble a bipolar spindle or to segregate chromosomes, but rather reflect a more specific requirement for CLASP/CLS-2.

A specific requirement for CLS-2 is further supported by our analysis of oocyte meiotic contractile ring assembly and dynamics in *katanin/mei-1* oocytes, which assemble spindles that lack any polarity and completely fail to organize the dispersed oocyte chromosomes throughout meiosis I (34, 39, 40). Furthermore, similar to *cls-2* oocytes, microtubule levels are substantially reduced in *mei-1* mutant oocytes (47). Nevertheless, stable contractile rings usually formed in *mei-1* mutant oocytes, although the rings appeared somewhat larger in diameter than in control oocytes, and we frequently observed extensive furrow ingressions that often enclosed the oocyte chromosomes but usually failed to complete constriction and regressed late in cytokinesis. We conclude that contractile rings can assemble and remain stable until late in polar body extrusion, even when oocyte meiotic spindle assembly is at least as severely defective as in *cls-2* mutant oocytes.

### Roles for CLS-2 and MEI-1 in contractile ring assembly and constriction

While our analysis indicates that CLS-2 is required early for the assembly or stability of the oocyte meiosis I contractile ring, we do not know if the defects reflect a direct or indirect requirement, or how CLS-2 functions at a molecular level to promote polar body extrusion. Consistent with studies showing that CLASP family members associate with microtubules and promote microtubule stability (22–25), *cls-2* mutant oocytes have reduced levels of spindle microtubules throughout meiosis I. Moreover, *cls-2* mutant oocytes fail to assemble bipolar spindles or segregate chromosomes, with ASPM-1 pole foci and chromosomes instead converging into a small amorphous cluster. While the contractile ring assembly defects we observed could be an indirect consequence of aberrant spindle assembly and structure, the more normal contractile ring dynamics observed in both *klp-18* and *mei-1* mutant oocytes suggest that CLS-2 has a more specific role in mediating ring assembly or stability. Notably, human CLASP proteins have been shown to associate with actin filaments and may cross-link microtubules and filamentous actin (26). Furthermore, mutations in the *Drosophila* CLS-2 ortholog Orbit/Mast cause spermatocyte cytokinesis defects that appear to result from a loss of central spindle microtubules that normally promote assembly of the contractile ring (48). Thus, it is possible that the contractile ring assembly or stability defects in *cls-2* mutants are due to a loss of microtubule and microfilament interactions that require CLS-2.

Consistent with a role for CLS-2 in bridging spindle microtubules and cortical actin filaments, we observed CLS-2∷GFP throughout the oocyte spindles at the time when contractile ring assembly begins. However, at the time that ring assembly begins, very few spindle microtubules have been observed in close proximity to the cell cortex (49). Biochemical studies of CLS-2 and its potential interactions with microtubules and actin filaments, along with higher resolution imaging of CLS-2, spindle microtubules and cortical microfilaments in oocytes may improve our understanding of how CLS-2 promotes the proper assembly and stability of the contractile ring during polar body extrusion.

Contractile ring dynamics during meiosis I were more normal after *mei-1* RNAi knockdown, but the furrows nevertheless often regressed and polar body extrusion usually failed. Previous studies have shown that partial loss of function mutations in *mei-1* result in abnormally large second polar bodies that are produced after meiosis II, as a result of decreased microtubule severing that is required for complete disassembly of the oocyte meiosis II spindle (42, 43). Our results provide the first systematic examination of polar body extrusion during meiosis I after depletion of katanin/MEI-1, and it is possible that the late failures in polar body cytokinesis that we observed also reflect a requirement for microtubule severing. Alternatively, similar defects are observed in oocytes lacking the centralspindlin components CYK-4 and ZEN-4 (10, 14, 27, 45), and it is possible that central spindle proteins are mis-regulated after MEI-1 knockdown. Further investigation of MEI-1 and its interactions with central spindle proteins may improve our understanding of this late requirement for MEI-1 during polar body extrusion.

### Negative regulation of oocyte global cortical dynamics and its relationship to polar body extrusion

In addition to the defects in contractile ring dynamics during polar body extrusion in *cls-2* mutant oocytes, we also observed increased membrane ingressions throughout the oocyte cortex. Furthermore, we detected CLS-2∷GFP in small patches throughout the cortex in control oocytes. These patches were present early in meiosis I but dissipated before anaphase chromosome segregation, prior to initiation of the membrane ingressions that occur during anaphase in wild-type oocytes. While the excessive global furrowing in *cls-2* mutants may result from loss of the cortical CLS-2 patches, if so then CLS-2 may act prior to the initiation of global cortical furrowing to down-regulate the ingressions. Alternatively, low levels of cortical CLS-2 that were not detected with our imaging methods might contribute to this negative regulation of membrane furrowing throughout the oocyte cortex.

The molecular mechanism by which cortical CLS-2 patches might down-regulate global membrane ingression during oocyte meiosis I is not clear. Our observations that NMY-2 and ANI-1 dynamics were not altered in *cls-*2 oocytes suggests that the increased membrane ingression is not be due to an increase in cortical actomyosin contractility but perhaps is an indirect consequence of defects in the stiffness of the oocyte cortex. The observations that CLASP orthologs can interact with both cortical microtubules and actin filaments may be consistent with such a role for CLS-2 (26). Moreover, oocyte polar body extrusion has been described as a bleb-like process, with local weakening of the actomyosin cytoskeleton inside the contractile ring promoting an out-pocketing of the membrane to form a polar body (5, 10). Studies of bleb formation in other cellular contexts have shown that microtubule destabilization can result in bleb formation (50–52). Thus, it is possible that loss of CLS-2 promotes membrane ingressions throughout the cortex due to destabilization of cortical microtubules and a corresponding decrease in cortical stiffness, although we have not yet detected statistically significant differences in the levels of cortical microtubules in *cls-2* oocytes (data not shown). Alternatively, mammalian CLASPs mediate linkage between microtubule plus ends and the cell cortex (53, 54), and such interactions might also influence cortical stiffness in *C. elegans* oocytes. Other factors, including the kinesin-13 family member KLP-7 and the TOG domain protein and XMAP215 ortholog ZYG-9, have been shown to down-regulate cortical microtubule levels during oocyte meiotic cell division in *C. elegans* (21, 55). Further investigation both of microtubule and actomyosin dynamics, using higher resolution light microscopy methods, and studies of how microfilament and microtubule regulators interact to influence oocyte cortical dynamics, may improve our understanding of how global cortical contractility and membrane dynamics influence contractile ring dynamics during polar body extrusion.

Comparing the consequences of reducing the functions of the casein kinase CNSK-1 and CLS-2 indicates that the relationship between global cortical dynamics and polar body extrusion is complex. CSNK-1 also limits membrane ingressions throughout the oocyte cortex during meiosis I but appears to do so through negative regulation of actomyosin dynamics, and CSNK-1 knockdown often results in extrusion of the entire meiotic spindle and all of the oocyte chromosomes into the first polar body (15). This outcome has led to a model in which global cortical contraction generates a hydrostatic cytoplasmic force that promotes an out-pocketing of the plasma membrane as the spindle is pushed through the contractile ring and into the forming polar body. While such a mechanism may operate, it also is clear that the contractile ring and associated plasma membrane ingress substantially prior to constricting roughly midway along the axis of the spindle during polar body extrusion. These dynamics suggest that the spindle and the contractile ring interact to promote furrow ingression and constriction. Moreover, the increased global membrane ingressions in *cls-2* oocytes do not appear to result from increased actomyosin contractility and are accompanied by defects in ring assembly or stability and a frequent failure to extrude a polar body. Thus, it appears that negative regulation of global cortical membrane ingression during oocyte meiotic cell division can both promote and prevent the extrusion of chromosomes into polar bodies. These different outcomes presumably reflect differences in how CSNK-1 and CLS-2 influence the cortical cytoskeleton and its dynamics. One possible explanation for these different outcomes is that a loss of cortical stiffness in *cls-2* oocytes might allow for increased membrane ingression throughout the oocyte cortex, while at the same time disrupting contractile ring assembly and/or preventing any increase in the hydrostatic cytoplasmic force that has been proposed to push the meiotic spindle partly into the budding polar body (15). Further investigation of cortical cytoskeleton dynamics, and the interactions of factors that regulate these dynamics during polar body extrusion, should improve our understanding of this poorly understood but fundamentally important biological process.

## Materials and methods

### *C. elegans* Strain Maintenance

All *C. elegans* strains used in this study (S1 Table), were maintained at 20°C as described previously (56).

### *cls-2* CRISPR/Cas9 Allele Generation

Mutations in *cls-2* were generated by injecting young adult N2 hermaphrodites with the following mixture (57): 25μM *cls-2* crRNA (ATCAGCCGATCGACTCCGGG), 5μM *dpy-10* crRNA (GCTACCATAGGCACCAC GAG), 30μM trRNA, 2.1μg/μl Cas9-NLS, and 2.5μM *dpy-10* single-strand DNA (ssDNA, CACTTGAACTTCAATACGGCAAGATGAGAATGACTGGAAACCGTACCGCATGCGGTGCCTATGGTAGCGGAGCTTCACATGGCTTCAGACCAACAGCCTAT). No homologous repair template was used for *cls-2,* and *cls-2* DNA breaks were allowed to repair randomly. Before injection, the trRNA and crRNAs were mixed and incubated at 95°C for 5 minutes, before cooling at room temperature for 5 minutes. After cooling, Cas9-NLS (QB3-Berkeley MacroLab) was added to the annealed trRNA and crRNAs and allowed to incubate for another 5 minutes at room temperature before the *dpy-10* ssDNA repair template was added. After injection, hermaphrodites were singled out and their broods were screened for *dpy-10* roller or dumpy co-conversion worms, which were allowed to produce broods. Those broods were then evaluated for potential *cls-2* phenotypes (embryonic lethality), and lines identified as potentially carrying mutations to *cls-2* were balanced. PCR amplified fragments were Sanger sequenced to identify the CRISPR/Cas9-induced mutations.

### Feeding RNAi Knockdown of *mei-1* and *klp-18*

RNAi knockdown of *mei-1* and *klp-18* was achieved by plating hypochlorite synchronized L1 larvae onto *E. coli* (HT115) lawns induced to express dsRNA corresponding to *mei-1* or *klp-18 (58)*. Plated worms were either maintained at 20°C until adults were imaged (*mei-1*) or maintained at 20°C and upshifted to 26°C 16 hours prior to imaging (*klp-18*) to ensure robust knockdown, as determined by the formation of monopolar meiotic spindles in both meiosis I and II. The *mei-1* RNAi vector was from the Ahringer RNAi library (59). The *klp-18* RNAi vector was made by amplifying a portion of the *klp-18* coding sequence from isolated N2 genomic DNA (using primers 5’-ACCGGCAGATCTGATATCATCGATGAATTCTCCAACTTTCAA ATGCCACA-3’ and 5’-ACGGTATCGATAAGCTTGATATCGAATTCCTTCGATATGGAA GAA AGCGG-3’), which was inserted into the L4440 vector backbone using the NEBuilder HiFi DNA assembly cloning kit (NEB).

### Live-cell Imaging

All imaging was carried out using a Leica DMi8 microscope outfitted with a spinning disk confocal unit - CSU-W1 (Yokogawa) with Borealis (Andor), dual iXon Ultra 897 (Andor) cameras, and a 100x HCX PL APO 1.4-0.70NA oil objective lens (Leica). Metamorph (Molecular Devices) imaging software was used for controlling image acquisition. The 488nm and 561nm channels were imaged simultaneously every 10 seconds with 1μm Z-spacing (either 16μm or 21μm total Z-stacks depending on the fluorescent markers used, with the same stack size used for all movies utilizing the same fluorescent markers).

*In utero* live imaging of oocytes was accomplished by mounting adult worms with a single row or less of embryos in 1.5μl of M9 mixed with 1.5μl of 0.1μm polystyrene Microspheres (Polysciences Inc.) on a 6% agarose pad with a coverslip gently laid over top. *Ex utero* imaging of oocytes was carried out by cutting open adult worms with a single row or less of embryos in 4μl of egg buffer (118mM NaCl, 48mM KCl, 2mM CaCl_2_, 2mM MgCl_2_, and 0.025 mM of HEPES, filter sterilized before HEPES addition) on a coverslip before mounting onto a 2% agarose pad on a microscope slide.

### Image analysis, Quantification, and Statistical Analysis

General image analysis and quantification of microtubules and global cortical furrowing was carried out using FIJI software (60). Three-dimensional projection and rotation of movies used to look at polar body contractile rings was carried out using Imaris software (Bitplane). Meiosis I polar body extrusion success was evaluated based on whether oocytes extruded any chromosomes marked by GFP or mCherry histone 2B (H2B) into a polar body that remained extruded for the period of imaging, either until meiosis I had obviously ended and meiosis II spindle assembly began, or until pronuclei began to decondense in the one-cell stage embryo after meiosis II. The end of meiosis I and beginning of meiosis II was considered to be the time at which the chromosomes left in the oocyte cytoplasm began to visibly separate from each other. Projections for spindle-associated furrow examination were made by manually isolating the 5 most spindle-associated z-planes for each time point during the period of global cortical furrowing and then sum projecting the mCherry∷PH membrane signal. Membrane temporal overlays were created by overlaying the outlined membrane regions of interest for the period of furrowing (detailed below) to create a single image.

Total spindle microtubule pixel intensity was determined using the following formula: (Mean Grey Value (spindle)/Mean Grey Value (cytoplasm)) × spindle area = total spindle microtubule pixel intensity. The mean grey values for both the meiotic spindle and cytoplasm were determined by drawing a region of interest around either the meiotic spindle or a portion of oocyte cytoplasm devoid of adjacent sperm in maximum projected Z-stacks and measuring the mean grey value of the selected region in ImageJ. Spindle area was determined by measuring the area of the region of interest encompassing the meiotic spindle.

Quantification of global cortical furrowing was accomplished by drawing regions of interest over the oocyte membrane signal (mCherry∷PH) for a single central z-slice for the entire period of global cortical furrowing. Regions of interest were then converted to a high contrast stack of membrane positions over time, which were then analyzed using the ADAPT plugin (61) for ImageJ in order to determine curvature values across the oocyte membrane. A furrow was defined as being at least two consecutive membrane points with negative mean curvature values and a standard deviation of mean curvature at least two standard deviations above the average standard deviation of mean curvature value for the entire oocyte membrane. Membrane points fitting the criteria of a furrow (above) that were separated by a single membrane point not fitting the criteria were considered as part of the same furrow for the purposes of counting. For statistical analysis of global cortical furrowing (Fig 7B), one-way ANOVA was used to determine if there was any difference in the mean furrowing between genotypes, F-Tests to compare the variances, and two-tailed Student’s t-Tests between genotypes to compare the means directly (assuming either equal or unequal variances depending on the F-Test results). Statistical analysis and graphs were completed using Excel (Microsoft).

## Acknowledgements

We thank Julien Dumont, Amy Maddox, and the Caenorhabditis Genetics Center (funded by the NIH Office of Research Infrastructure Programs; P40 OD010440) for *C. elegans* strains, Jie Yang for the KLP-18 RNAi vector, Chris Doe and Diana Libuda for sharing laboratory equipment, and members of the Bowerman laboratory for helpful discussions.

---

**S1 Fig - CLS-2 localizes to meiotic spindles and is required for their assembly**

(A) *In utero* time-lapse spinning disk confocal images of CLS-2∷GFP and mCherry∷H2B. (B) Protein domain maps of wild type CLS-2 and CRISPR-generated *cls-2* alleles *or1949*, *or1950*, and *or1951*. Each mutation results in multiple early stop codons before the first TOG domain, with the first stop codon indicated. (C) *In utero* time-lapse spinning disk confocal images of control and *cls-2* mutant oocytes with GFP∷TBB-2 and mCherry∷H2B. t = 0 seconds corresponds to nuclear envelope breakdown.

**S2 Fig - Control oocyte spindle-associated membrane furrows**

Time-lapse spinning disk confocal images of control oocytes expressing mCherry∷PH and GFP∷H2B; t = 0 seconds here and in subsequent Fig 4 related supplements (S3-6 Figs) corresponds to the time point immediately before global cortical furrowing begins, unless otherwise stated.

**S3 Fig - *cls-2(or1948)* oocyte spindle-associated membrane furrows**

Time-lapse spinning disk confocal images of *cls-2* mutant oocytes expressing mCherry∷PH and GFP∷H2B.

**S4 Fig - *cls-2(or1948)* oocyte spindle-associated membrane furrows**

Time-lapse spinning disk confocal images of *cls-2* mutant oocytes expressing mCherry∷PH and GFP∷H2B.

**S5 Fig - *klp-18(RNAi)* oocyte spindle-associated membrane furrows**

Time-lapse spinning disk confocal images of *klp-18(RNAi)* oocytes expressing mCherry∷PH and GFP∷H2B.

**S6 Fig - *mei-1(RNAi)* oocyte spindle-associated membrane furrows**

Time-lapse spinning disk confocal images of *mei-1(RNAi)* oocytes expressing mCherry∷PH and GFP∷H2B.

**S7 Fig - Control oocyte NMY-2∷GFP contractile rings**

Three-dimensionally projected and rotated spinning disk confocal time-lapse images of control oocytes expressing NMY-2∷GFP and mCherry∷H2B; t = 0 seconds in this and subsequent Fig 5 related supplements (S8-10 Figs) corresponds to the end of meiosis I and beginning of meiosis II (see Materials and methods).

**S8 Fig - *cls-2(or1948)* oocyte NMY-2∷GFP contractile rings**

Three-dimensionally projected and rotated spinning disk confocal time-lapse images of *cls-2* mutant oocytes expressing NMY-2∷GFP and mCherry∷H2B.

**S9 Fig - *klp-18(RNAi)* oocyte NMY-2∷GFP contractile rings**

Three-dimensionally projected and rotated spinning disk confocal time-lapse images of *klp-18(RNAi)* oocytes expressing NMY-2∷GFP and mCherry∷H2B.

**S10 Fig - *mei-1(RNAi)* oocyte NMY-2∷GFP contractile rings**

Three-dimensionally projected and rotated spinning disk confocal time-lapse images of *mei-1(RNAi)* oocytes expressing NMY-2∷GFP and mCherry∷H2B.

**S11 Fig - Control oocyte membrane temporal overlays**

Control oocyte membrane temporal overlays depicting membrane positions over time at a single focal plane throughout meiosis I.

**S12 Fig - *cls-2(or1948)* oocyte membrane temporal overlays**

*cls-2* mutant oocyte membrane temporal overlays depicting membrane positions over time at a single focal plane throughout meiosis I. Asterisks indicate oocytes in which polar body extrusion failed.

**S13 Fig - *klp-18(RNAi)* and *mei-1(RNAi)* membrane temporal overlays**

*klp-18(RNAi) and mei-1(RNAi)* oocyte membrane temporal overlays depicting membrane positions over time at a single focal plane throughout meiosis I. Asterisks indicate oocytes in which polar body extrusion failed.

**S14 Fig - Control oocyte NMY-2∷GFP cortical dynamics**

Time-lapse spinning disk confocal images of control oocytes expressing NMY-2∷GFP and mCherry:H2B; t = 0 seconds corresponds to the end of meiosis I and beginning of meiosis II in this and subsequent Fig 7 related supplements (S15-18 Figs).

**S15 Fig - Control oocyte mNG∷ANI-1 cortical dynamics**

Time-lapse spinning disk confocal images of control oocytes expressing mNeonGreen∷ANI-1 and mCherry∷H2B.

**S16 Fig - *cls-2(or1948)* oocyte NMY-2∷GFP cortical dynamics**

Time-lapse spinning disk confocal images of *cls-2* mutant oocytes expressing NMY-2∷GFP and mCherry∷H2B. All oocytes shown succeeded in polar body extrusion.

**S17 Fig - *cls-2(or1948)* oocyte NMY-2∷GFP cortical dynamics**

Time-lapse spinning disk confocal images of *cls-2* mutant oocytes expressing NMY-2∷GFP and mCherry∷H2B. All oocytes shown failed in polar body extrusion.

**S18 Fig - *cls-2(or1948)* oocyte mNG∷ANI-1 cortical dynamics**

Time-lapse spinning disk confocal images of *cls-2* mutant oocytes expressing mNeonGreen∷ANI-1 and mCherry∷H2B; t = 0s corresponds to the end of meiosis I and beginning of meiosis II. All oocytes shown failed in polar body extrusion.

**S1 Movie - *Ex utero* CLS-2∷GFP localization**

*Ex utero* time-lapse spinning disk confocal movie of a maximum projected oocyte expressing CLS-2∷GFP (green) and mCherry∷H2B (magenta). Frame rate is 10 frames per second.

**S2 Movie - *In utero* CLS-2∷GFP localization**

*In utero* time-lapse spinning disk confocal movie of a maximum projected oocyte expressing CLS-2∷GFP (green) and mCherry∷H2B (magenta). Frame rate is 10 frames per second.

**S3 Movie - Control oocyte membrane furrowing**

*Ex utero* time-lapse spinning disk confocal movie of a control oocyte expressing mCherry∷PH (black) and GFP∷H2B (magenta). In this and subsequent oocyte membrane furrowing videos, the 5 focal planes that encompassed most of the meiotic chromosomes were used; membrane images were sum projected, histones images were maximum projected. In this and all subsequent Fig 4 related movies (S4-10 Movies), the frame rate is 5 frames per second.

**S4 Movie - Control oocyte membrane furrowing**

*Ex utero* time-lapse spinning disk confocal movie of a control oocyte expressing mCherry∷PH (black) and GFP∷H2B (magenta).

**S5 Movie - *cls-2(or1948)* oocyte membrane furrowing**

*Ex utero* time-lapse spinning disk confocal movie of a *cls-2(or1948)* oocyte expressing mCherry∷PH (black) and GFP∷H2B (magenta).

**S6 Movie - *cls-2(or1948)* oocyte membrane furrowing**

*Ex utero* time-lapse spinning disk confocal movie of a *cls-2(or1948)* oocyte expressing mCherry∷PH (black) and GFP∷H2B (magenta).

**S7 Movie - *klp-18(RNAi)* oocyte membrane furrowing**

*Ex utero* time-lapse spinning disk confocal movie of a *klp-18(RNAi)* oocyte expressing mCherry∷PH (black) and GFP∷H2B (magenta).

**S8 Movie - *klp-18(RNAi)* oocyte membrane furrowing**

*Ex utero* time-lapse spinning disk confocal movie of a *klp-18(RNAi)* oocyte expressing mCherry∷PH (black) and GFP∷H2B (magenta).

**S9 Movie - *mei-1(RNAi)* oocyte membrane furrowing**

*Ex utero* time-lapse spinning disk confocal movie of a *mei-1(RNAi)* oocyte expressing mCherry∷PH (black) and GFP∷H2B (magenta).

**S10 Movie - *mei-1(RNAi)* oocyte membrane furrowing**

*Ex utero* time-lapse spinning disk confocal movie of a *mei-1(RNAi)* oocyte expressing mCherry∷PH (black) and GFP∷H2B (magenta).

**S11 Movie - Control oocyte NMY-2∷GFP contractile ring dynamics**

*Ex utero* 3-dimensionally projected and rotated time-lapse spinning disk confocal movie of control oocyte expressing NMY-2∷GFP (green) and mCherry:H2B (magenta). In this and all subsequent Fig 5 related movies (S12-18 Movies), the frame rate is 5 frames per second.

**S12 Movie - Control oocyte NMY-2∷GFP contractile ring dynamics**

*Ex utero* 3-dimensionally projected and rotated time-lapse spinning disk confocal movie of control oocyte expressing NMY-2∷GFP (green) and mCherry:H2B (magenta).

**S13 Movie - *cls-2(or1948)* oocyte NMY-2∷GFP contractile ring dynamics**

*Ex utero* 3-dimensionally projected and rotated time-lapse spinning disk confocal movie of *cls-2(or1948)* oocyte expressing NMY-2∷GFP (green) and mCherry:H2B (magenta).

**S14 Movie - *cls-2(or1948)* oocyte NMY-2∷GFP contractile ring dynamics**

*Ex utero* 3-dimensionally projected and rotated time-lapse spinning disk confocal movie of *cls-2(or1948)* oocyte expressing NMY-2∷GFP (green) and mCherry:H2B (magenta).

**S15 Movie - *klp-18(RNAi)* oocyte NMY-2∷GFP contractile ring dynamics**

*Ex utero* 3-dimensionally projected and rotated time-lapse spinning disk confocal movie of *klp-18(RNAi)* oocyte expressing NMY-2∷GFP (green) and mCherry:H2B (magenta).

**S16 Movie - *klp-18(RNAi)* oocyte NMY-2∷GFP contractile ring dynamics**

*Ex utero* 3-dimensionally projected and rotated time-lapse spinning disk confocal movie of *klp-18(RNAi)* oocyte expressing NMY-2∷GFP (green) and mCherry:H2B (magenta).

**S17 Movie - *mei-1(RNAi)* oocyte NMY-2∷GFP contractile ring dynamics**

*Ex utero* 3-dimensionally projected and rotated time-lapse spinning disk confocal movie of *mei-1(RNAi)* oocyte expressing NMY-2∷GFP (green) and mCherry:H2B (magenta).

**S18 Movie - *mei-1(RNAi)* oocyte NMY-2∷GFP contractile ring dynamics**

*Ex utero* 3-dimensionally projected and rotated time-lapse spinning disk confocal movie of *mei-1(RNAi)* oocyte expressing NMY-2∷GFP (green) and mCherry:H2B (magenta).

**S19 Movie - Control oocyte mNG∷ANI-1 contractile ring dynamics**

*Ex utero* 3-dimensionally projected and rotated time-lapse spinning disk confocal movie of control oocyte expressing mNG∷ANI-1 (green) and mCherry:H2B (magenta). In this and all subsequent Fig 6 related movies (S20-22 Movies), the frame rate is 5 frames per second.

**S20 Movie - *cls-2(or1948)* oocyte mNG∷ANI-1 contractile ring dynamics**

*Ex utero* 3-dimensionally projected and rotated time-lapse spinning disk confocal movie of *cls-2(or1948)* oocyte expressing mNG∷ANI-1 (green) and mCherry:H2B (magenta).

**S21 Movie - Control oocyte NMY-2∷mKate2 and mNG∷ANI-1 contractile ring dynamics**

*Ex utero* 3-dimensionally projected and rotated time-lapse spinning disk confocal movie of control oocyte expressing NMY-2∷mKate2 (magenta), mNG∷ANI-1 (green), and mCherry:H2B (magenta).

**S22 Movie - *cls-2(or1948)* oocyte NMY-2∷mKate2 and mNG∷ANI-1 contractile ring dynamics**

*Ex utero* 3-dimensionally projected and rotated time-lapse spinning disk confocal movie of *cls-2(or1948)* oocyte expressing NMY-2∷mKate2 (magenta), mNG∷ANI-1 (green), and mCherry:H2B (magenta).

**S23 Movie - Control oocyte NMY-2∷GFP cortical dynamics**

Three example time-lapse spinning disk confocal movies of *ex utero* control oocytes expressing NMY-2∷GFP (green) and mCherry∷H2B (magenta). In this and all subsequent Fig 7 related movies (S24-26 Movies), the frame rate is 5 frames per

**S24 Movie - *cls-2(or1948)* oocyte NMY-2∷GFP cortical dynamics**

Three example time-lapse spinning disk confocal movies of *ex utero cls-2(or1948)* oocytes expressing NMY-2∷GFP (green) and mCherry∷H2B (magenta).

**S25 Movie - Control oocyte mNG∷ANI-1 cortical dynamics**

Three example time-lapse spinning disk confocal movies of *ex utero* control oocytes expressing mNG∷ANI-1 (green) and mCherry∷H2B (magenta).

**S26 Movie - *cls-2(or1948)* oocyte mNG∷ANI-1 cortical dynamics**

Three example time-lapse spinning disk confocal movies of *ex utero cls-2(or1948)* oocytes expressing mNG∷ANI-1 (green) and mCherry∷H2B (magenta).

**S1 Table - Table of *C. elegans* strains used in this study**

